# ISG15 Monomer Promotes IFNα-mediated Antiviral Activity against Pseudorabies Virus by Facilitating phosphorylation of STAT1/STAT2

**DOI:** 10.1101/2022.05.02.490374

**Authors:** Huimin Liu, Chen Li, Wenfeng He, Jing Chen, Guoqing Yang, Lu Chen, Hongtao Chang

## Abstract

Pseudorabies virus (PRV), which presently lacks both an antiviral drug and a viable therapeutic option, is a major viral pathogen that poses a danger to the pig industry worldwide. Interferon-stimulated gene 15 (ISG15) is strongly upregulated during viral infections and has been reported to have proviral or antiviral properties, depending on the virus and host species. Our previous studies demonstrated ISG15 was remarkably upregulated during PRV infection, and the overexpression of ISG15 inhibited PRV replication. Nevertheless, the exact mechanism through which ISG15 influences PRV replication poorly understood unclear. Here, we demonstrate that ISG15 accumulation induced by PRV infection requires viral gene expression and viral growth. Conjugation inhibition assays showed that ISG15 imposes its antiviral effects via unconjugated (free) ISG15 and affects the viral release. In addition, we found ISG15 promoted IFN*α*-mediated antiviral activity against PRV by facilitating the phosphorylation of STAT1 and STAT2, along with an increase of ISGF3-induced ISRE promoter activity. Furthermore, we evaluated the role of ISG15 in host defense to control PRV infection by using ISG15^-/-^ mice model. When challenged with PRV, ISG15^-/-^ mice exhibited increased morbidity and mortality, as well as viral load compared to WT mice. Pathological examination revealed increased lesions, mononuclear cellular infiltration and neuronal death in the brains of ISG15^-/-^ mice, along with the upregulation of the cytokines. Our findings establish the importance of ISG15 in IFN*α*-induced antiviral response and in the control of PRV infection.

## Introduction

Pseudorabies virus (PRV), an alphaherpesviruses, is a significant viral pathogen of pigs, wreaking havoc on the global pig industry [1]. PRV is able to establish persistent infection in peripheral neurons of the host with no specific clinical symptoms, which is usually as a useful model for understanding alpha herpes virus biology and host’s innate immune response [2, 3]. Recent evidence indicates that PRV can induce severe clinical symptoms in people, such as acute encephalitis, under specific conditions [4-8]. Despite intensive research, neither specific antiviral therapy nor effective vaccines against PRV are currently available [4, 9]. Therefore, a better understanding of the interactions between PRV infection and the host responses that inhibit PRV infection is of great importance.

In response to the viral invasion, the host evolves various defense mechanisms. Among these, the type I interferon (IFN-I) plays a central role in host defense against viral infections. IFN-I, represented by IFN*α* and IFN*β*, binds to their respective receptors and activates the JAKs, which subsequently phosphorylate STAT1 and STAT2. The phosphorylated (p-) STAT1 and p-STAT2 complex with IRF9, resulting in the formation of ISG factor 3 (ISGF3). ISGF3 shuttles to the nucleus, where it binds to the IFN-stimulated response element (ISRE) in DNA and stimulates the transcription of hundreds of interferon-stimulated genes (ISGs) involved in the host antiviral response [10, 11]. There is increasing evidence that PRV encodes proteins to antagonize the IFN response by suppressing IFN production, blocking IFN downstream signaling, or regulating specific ISGs [12-14]. ISG15, a ubiquitin-like modifier, is among the most frequently induced proteins by IFN-I and has been reported to have proviral or antiviral activities, depending on the virus and host species [15]. Like ubiquitination, ISG15 may be covalently attached to substrates via a conserved C-terminal Gly-Gly motif. This process is termed ISGylation through a cascading reaction catalyzed by E1 activating (UbE1L), E2 conjugating (UbcH8), and E3 ligase enzymes (Herc5), which are also induced by IFN-I, which has been demonstrated to cause either a gain or a loss of function of target proteins [16]. ISG15 can be removed from its target proteins by the ubiquitin-specific protease USP18, making the ISGylation process reversible [17-20]. In addition to its conjugated form, unconjugated (free) ISG15 plays cytokine activity when released to the extracellular

With respect to PRV, our previous study showed that ISG15 is greatly upregulated during PRV infection, and ISG15 overexpression inhibits PRV replication [21]. However, the mechanism underlying the anti-PRV effect of ISG15 *in vitro* and *in vivo* remains unexplored. Here, we characterized ISG15 expression profiles during PRV infection, and found that ISG15 inhibits PRV replication via the ISG15 monomer. Significantly, ISG15 silencing impairs IFN*α*-mediated anti-PRV effect by blocking phosphorylation of STAT1 and STAT2. Furthermore, ISG15 knockout mice exhibited enhanced PRV infection, as evidenced by high mortality, increased viral titer and severe inflammatory. These results reveal a critical role for ISG15 in IFN*α*-induced host antiviral activity and will provide a potential cellular therapeutic strategy.

## RESULTS

### ISG15 monomer and conjugated protein accumulation during PRV infection

Although ISG15 expression has reportedly increased during PRV infection in our previous study, how free and conjugated ISG15 proteins impact PRV infection and their roles in host defenses against PRV infection remain largely unknown. We sought to dissect the ISG15 behavior during PRV infection by analyzing the mRNA levels of ISG15 and PRV glycoprotein E (gE), a late viral gene, at different times post infection. As shown in Figure 1A, PRV infection induces a progressive increase in ISG15 RNA in a time-dependent manner. The increase started at 6 hours post-infection (hpi), reaching the peak level at 24 hpi. PRV-gE RNA increase trend was consistent with ISG15 RNA (Fig.1A).

**FIG 1.**
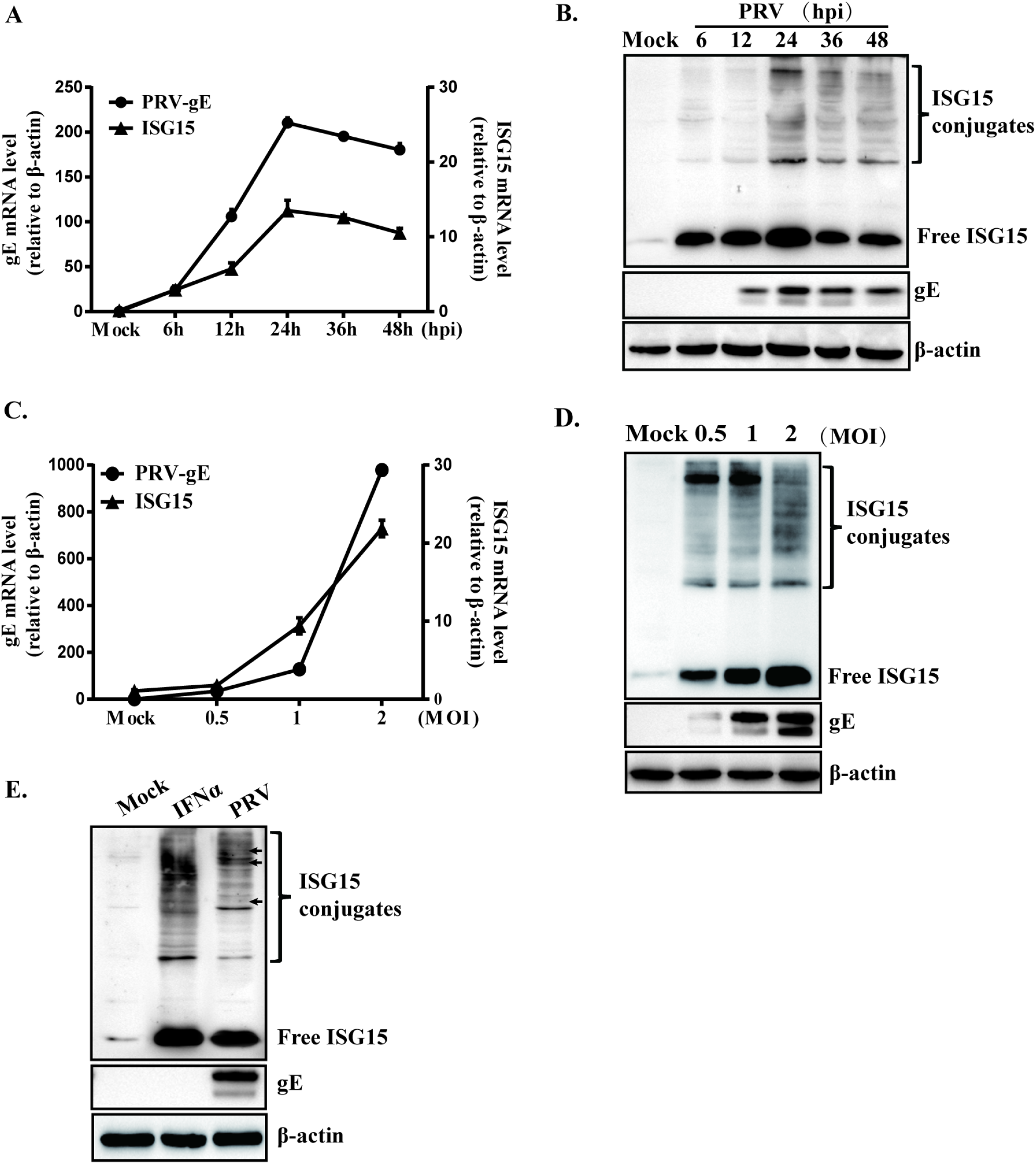
Upregulation of ISG15 and ISGylation expression during PRV infection. (A) PK15 cells were infected with PRV (MOI=1), and supernatants were collected at various time points following infection. PRV-gE and ISG15 mRNA levels were detected by RT-qPCR. The data represent the fold increase in gE and ISG15 RNA levels in PRV-infected cells relative to mock-infected cells. (B) Total protein extracts from panel A were tested for immunoblot analysis using antibodies specific for ISG15, PRV-gE, or *β*-actin. (C) PK15 cells were either mock-infected or infected with PRV at indicated MOI, and total RNA was harvested 24 hpi. RT-qPCR was used to quantify PRV-gE and ISG15 mRNA level. (D) As in panel C, Western blot analysis was used to determine ISG15 expression and protein ISGylation accumulation. (E) ISG15 expression patterns induced by IFNα (1000 U/ml) or PRV infection were compared by Western blotting. ** p < 0.01 by Student’*s* test.

To confirm the results mentioned above, immunoblotting was used to determine if ISG15 and PRV-gE RNA levels were translated into protein levels. Accumulation of unconjugated ISG15 and elevated high-molecular-weight ISG15-conjugated proteins were detected from the above parallel infection samples. Unconjugated ISG15 monomer were detected beginning at 12 hpi, while the abundance of the ISG15-conjugated protein was observed at 6 hpi (Fig.1B). Meanwhile, PRV-gE expression started to be detected at 12 hpi and reached the maximum level at 24 hpi (Fig.1B). Compared to mock-infected cells, increased levels of ISG15-conjugated and free ISG15 proteins persisted over a 48-hour time period, reaching maximum level at 24 hpi. This finding suggested that the increase of ISG15 monomer/conjugates was related to an increase in PRV-gE abundance. To generate additional support for this point, we conducted a dose-response investigation by infecting PRV at different multiplicities of infection (MOI). We found that the overall ISG15 monomer and conjugation levels induced by PRV infection in a dose-dependent manner (Fig. 1C to 1D).

Finally, the ISG15 expression patterns induced by PRV infection or IFN stimulation were compared by Western blotting. The obtained data suggested that PRV-induced ISGylation differs to some extent from that of IFNα, as well as some specific bands were apparent in PRV-infected cells (Fig.1E).

These results suggested that ISG15 accumulation was induced early in the PRV replication cycle and began to enhance with the expression of late viral gene. Therefore, we speculate the ISG15 abundance may be associated with viral DNA synthesis or late viral gene expression.

### ISG15 accumulation are triggered by PRV gene expression

To analyze whether viral DNA synthesis impacts free and conjugated ISG15 accumulation, PK15-infected cell cultures were treated with phosphonoacetic acid (PAA) to suppress viral DNA synthesis and late gene expression. As expected, PAA effectively reduced the transcript and expression of ISG15, along with the inhibition of PRV-gE protein expression (Fig. 2A). Similar levels of unconjugated and conjugated ISG15 proteins were observed in PAA-treated cultures following mock- or infected with PRV. Thus, blocking late gene expression with PAA significantly decreased ISG15 accumulation.

**FIG 2.**
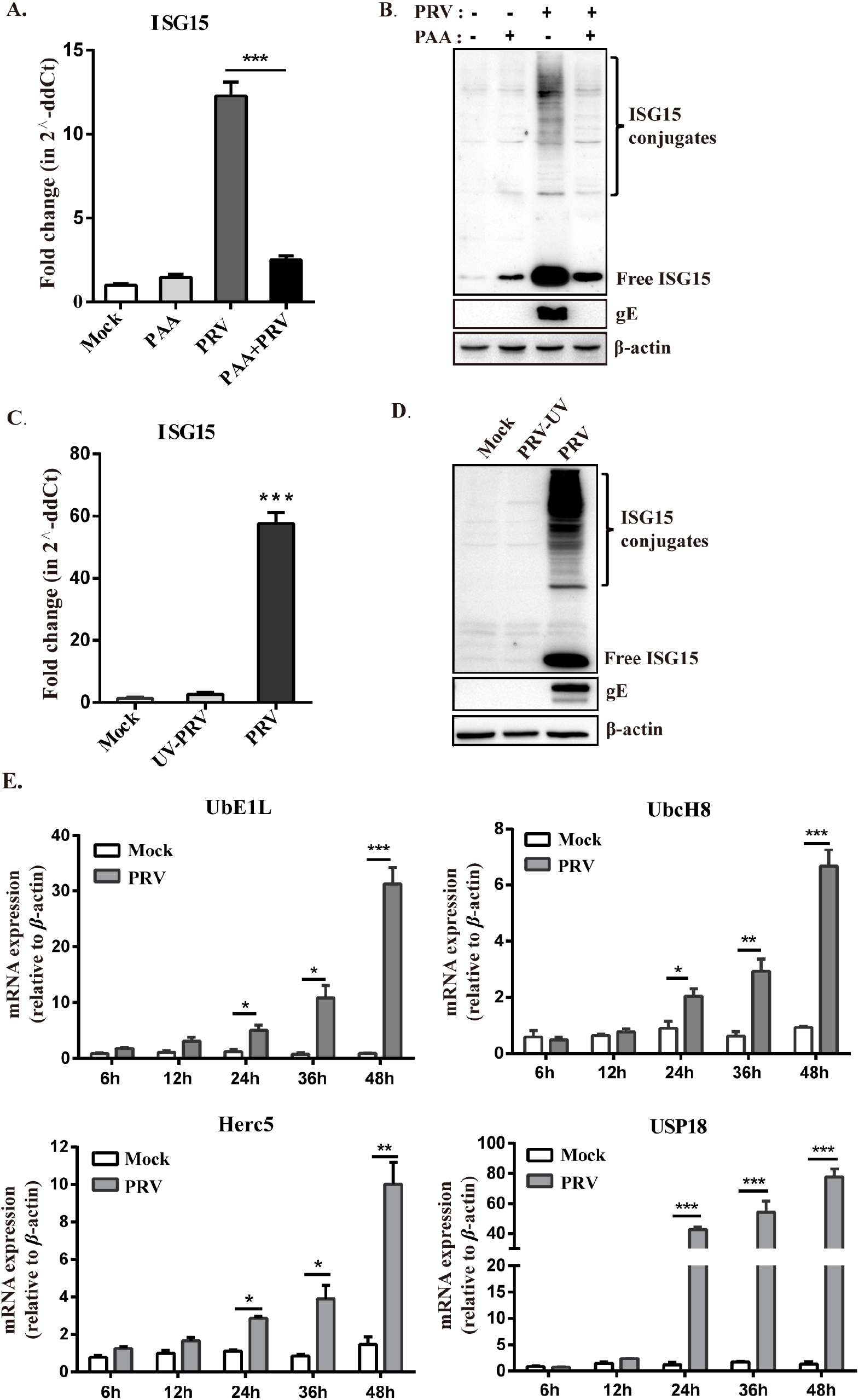
Regulation of free ISG15 and ISGylation accumulation in PRV-infected PK15 cells. (A and B) PK15 cells were mock-infected and PRV-infected left untreated or treated with PAA. Total RNA was collected 24 hpi and ISG15 mRNA was detected by RT-qPCR (A). Total proteins were harvested 24 hpi and the protein levels of ISG15 and PRV-gE were analyzed by Western blot (B). (C and D) The transcript and expression levels of ISG15 were detected in PK15 cells infected with PRV or UV-inactivated PRV 24 hpi, by RT-qPCR and Western blot respectively. (E) Total RNA was collected from PK15 cells mock-infected or PRV-infected for 24 h, and RT-qPCR was used to determine the mRNA levels of ISGylation enzymes (UbE1L, UbcH8, Herc5 and USP18). *, *p* < 0.05; **, *p* < 0.01; ***, *p* < 0.001 (*t*-test).

For further evidence, PRV virions were inactivated by UV irradiation [22], and viral inactivation was verified by a lack of viral growth after 24h post-inoculation. Infection of PK15 cells with UV-inactivated PRV resulted in greatly reduce in the transcription of ISG15 as well as free and conjugated ISG15 proteins levels, in comparison with that in cultures infected with active, unirradiated PRV (Fig. 2B). In addition, the conjugated ISG15 abundance is most likely via promoting an E1-E2-E3 enzymatic cascade of ISGylation. Overall mRNA levels of the UbE1L, UbcH8, Herc5 and USP18 progressively rised, demonstrating that the PRV replication process activates the ISGylation pathway (Fig. 2C). These results indicated that active PRV replication was needed for the free ISG15 and conjugate expression.

### ISG15 silencing enhances PRV replication

To analyze how ISG15 impacts PRV replication, ISG15 was deleted (ISG15^-/-^) in PK15 cells using CRISPER/Cas9 editing technology in our previous study [23]. ISG15 wild-type (WT) and ISG15^-/-^ cells were either mock-infected or infected with PRV at an MOI of 1. A significantly higher of amount of PRV-gE protein was observed in ISG15^-/-^ cells compared to WT cells at various time points post infection (Fig. 3A). Additionally, with the increase of MOI, the magnitude of this effect was also increased (Fig. 3A). Similar changes in PRV replication were found between WT and ISG15^-/-^ cells infected with PRV for 24h, as evidenced by fluorescence microscopy (Fig. 3B). Furthermore, a significant growth in PRV titer was detected in ISG15^-/-^ cells in comparison with WT cells at the same time point following infection (Fig.3C). Our findings established that ISG15 silencing accelerated PRV replication.

**FIG 3.**
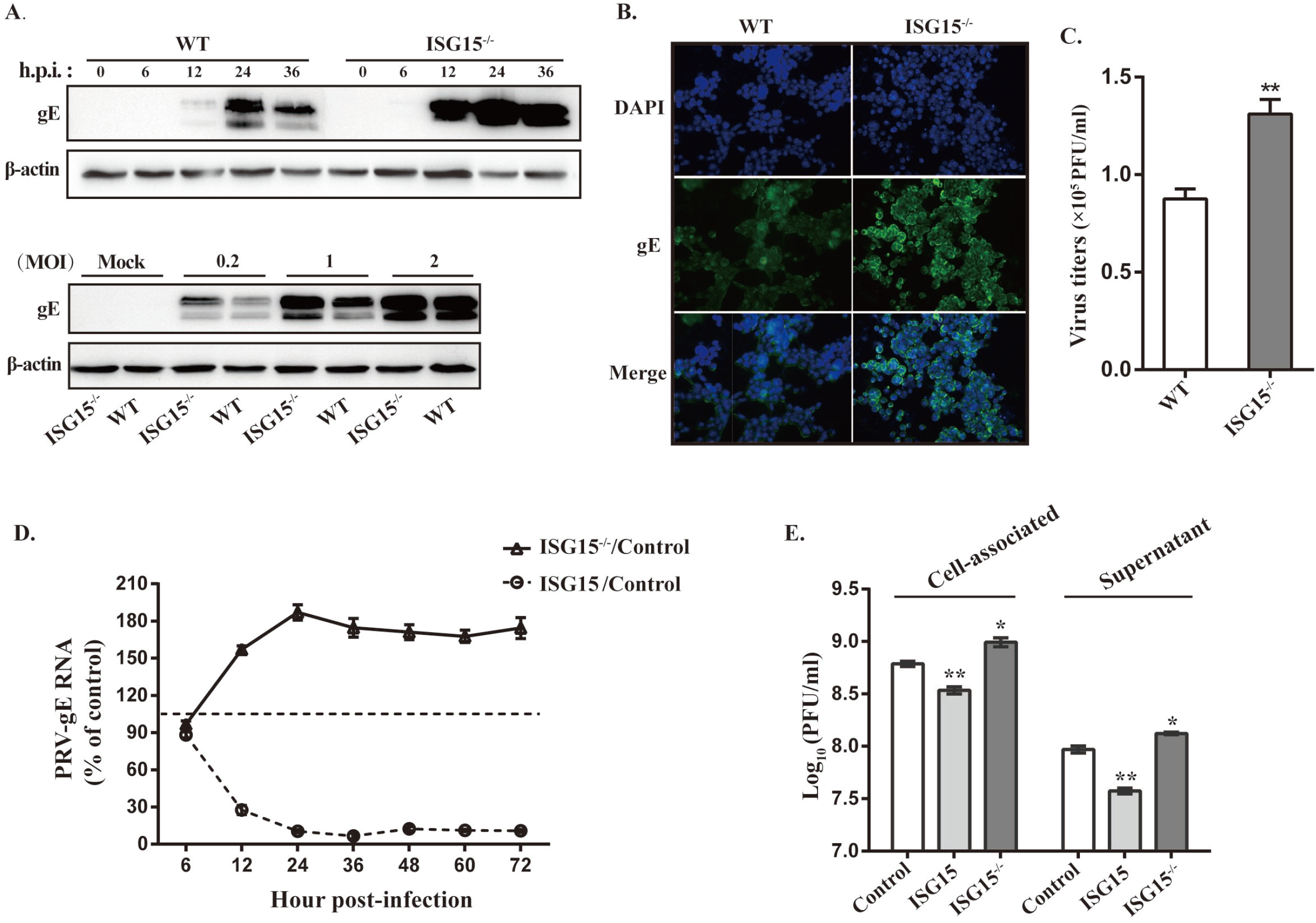
ISG15 inhibits PRV replication. (A and B) PRV-gE protein levels were detected in WT and ISG15^-/-^ cells at identified time points and MOI by Western blot. (C) WT and ISG15^-/-^ cells were infected with PRV (MOI=1) for 24 h, and the PRV-gE protein was measured using immunofluorescence microscopy assay. (D) PK15 cells were transfected with ISG15-overexpressing plasmid and then infected with PRV (MOI=1) 12 h later. PRV-gE RNA was quantified by RT-qPCR at different times post-infection. The data represent the percentages of expression of PRV-gE in ISG15-transfected cells or ISG15 knockout cells compared with cells transfected with a control plasmid (100%). (E) The PRV titer in the culture supernatant or associated with cells was determined by plaque assay at 24 hpi. Each experiment was repeated at least three times separately. *, p < 0.05; ** p < 0.01 (*t*-test).

It has been reported that ISG15 may affect virus entry [24] or release [25, 26], so we next sought to determine the role of ISG15 in those steps of PRV replication. We first determined whether ISG15 disrupts PRV replication by impacting virus entry into the cells. PK15 cells were transfected with a plasmid expressing ISG15 or a control plasmid before being infected with PRV, meanwhile, ISG15^-/-^ cells were also infected with PRV. Then, the PRV-gE RNA was quantified by RT-qPCR at various times post-infection. PRV-gE, a protein encoded by late gene, participates in virion assembly and release [27]. The results identified an obvious reduction of the PRV-gE RNA in the ISG15-overexpressing cells compared to control cells starting 6 hpi, while loss of ISG15 significantly enhance PRV-gE RNA level (Fig. 3D). This shows that ISG15 restricts PRV growth at a post-entry stage of infection. Additionally, the virus associated with cells and released to the supernatant were also detected by plaque assay. As illustrated in Figure 3E, a significant decrease was observed in virus titers from cell-associated fraction of WT cells expressing ISG15 compared to the same fraction of WT cells, while an increase in ISG15 knockout cells (Fig.3E, Cell-associated). Similarly, more than 3-fold reduction in virus titer was observed in the supernatant of WT cells overexpressing ISG15 with respect to WT and ISG15^-/-^ cells. This indicates that ISG15 limits PRV replication occurring before virus release.

Combinedly, these findings suggest that ISG15 generated by host cells inhibits PRV replication, possibly due to the virus release inhibition.

### Antiviral activity of ISG5 against PRV is due to ISG15 monomer

To further determine whether ISG15 achieves its antiviral effect against PRV via free or conjugated form, three different experiments were carried out.

First, due to the fact that exposure of the C-terminal Gly-Gly motif is essential for conjugation of ISG15 to substrates [28], these two residues were replaced with Ala using site-directed mutagenesis to generate a mutate plasmid ISG15AA employed as a control. PK15 cells were transfected with an emptor control vector, a ISG15-expressing plasmid, or an ISG15AA expressing plasmid. After 24 hours, the cells were infected with PRV and then viral protein expression and viral titer were measured. As expected, we found a significant decrease in PRV-gE expression level, as well as PRV titer (3.2-fold) (Fig. 4A), compared to cells transfected with empty vector. Notably, there wasn’t any significant differences detected in PRV titer and protein levels between these cells transfected with ISG15 and ISG15AA plasmid (Fig. 4A). Similar results were obtained from a parallel transfection / infection in ISG15^-/-^ cells (Fig. 4B). These data suggest that ISG15 exerts its antiviral activity against PRV through ISG15 monomer-dependent or conjugation-independent mechanisms.

**FIG 4.**
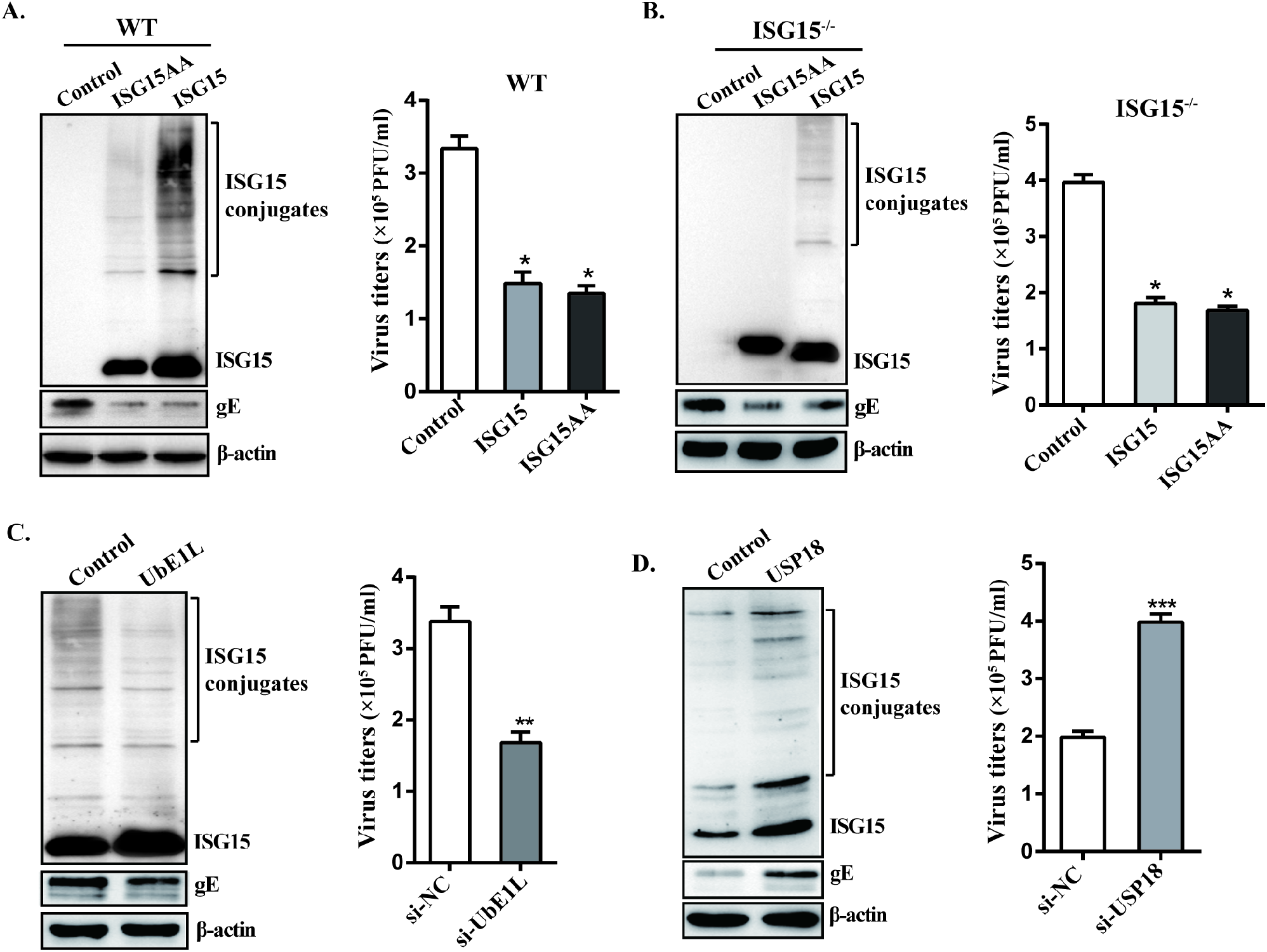
Antiviral activity of ISG15 against PRV relies on free ISG15. (A) PK15 cells were transfected with either an empty control plasmid, an ISG15-expressing plasmid, or an plasmid expressing ISG15 mutant (ISG15AA). After 24 hours, the cells were infected with PRV (MOI=1). Total proteins were collected 24 hpi, and Western blot was used to determine the expression of ISG15 unconjugated and conjugated protein. The PRV titer was detected by plaque assay. (B) As in panel A, ISG15^-/-^ cells were transfected and infected, and the ISG15 protein expression and virus titer were determined. (C to D) PK15 cells were transfected with either control siRNA or UbE1L siRNAs (C), USP18 siRNAs (D), and infected 24 h later with PRV (MOI=1). Protein extracts were harvested at 24 hpi and determined by Western blot with anti-ISG15 and anti-gE antibodies. Total RNA was collected to determine PRV titer by plaque assay. *, *p* < 0.05; **, *p* < 0.01; ***, *p* < 0.001 (*t*-test).

To verify these results and assess the influence of free ISG15 on PRV infection, a second approach was adopted by knocking down UbE1L to prevent the synthesis of ISG15 conjugates [29]. Prior to infection with PRV, PK15 cells were transfected with either a control siRNA or a siRNA targeting UbE1L, and ISG15 un-conjugated and conjugated proteins abundances were measured by immunoblotting. We observed that UbE1L-silenced cells displayed higher free ISG15 than control cells, nonetheless, a decrease in the expression levels of ISG15 conjugates (Fig. 4C). Furthermore, when UbE1L-silenced cells compared with the control cells, a significant drop in viral titer was found (Fig. 4C).

The third strategy involved silencing exogenous USP18, a deconjugating protease specific for ISG15 that removes ISG15 from targets [30]. This result is quite the opposite of the results from UbE1L-silencing experiments. Knockdown of USP18 resulted in an increased induction of the protein ISGylation, obvious enhancement in PRV-gE protein expression and viral production (Fig. 4D). This indicated that ISG15 inhibits PRV replication via free ISG15.

Collectively, these results indicate that free ISG15, rather than ISGylation, possesses an important antiviral activity against PRV.

### ISG15 contributes to IFN*α* antiviral activity against PRV

Previous reports have indicated that IFNα treatment of ISG15-deficient patient cells increased their resistance to several viral infection by viruses [31]. PRV is capable of establishing persistent infections, partly due to its ability to circumvent the host’s antiviral defenses, notably the type I IFN [13, 27]. Thus, to investigate whether ISG15 is involved in IFN-I mediated antiviral effect, WT and ISG15^-/-^ cells infected with PRV with or without IFN*α* treatment. Compared with their respective controls, the expression of PRV-gE and viral productions in ISG15^-/-^ cells were obviously increased when treated with IFN*α* (Fig.5A to 5C). This finding implies that complete loss of ISG15 impairs IFN*α* antiviral response against PRV.

To confirm this effect was caused by free ISG15, we performed gene rescue of ISG15 by transfecting a plasmid expressing ISG15AA in ISG15^-/-^ cells. A similar level of PRV-gE expression was detected in ISG15^-/-^ cells transfected with ISG15AA-expressing plasmid and WT cells, indicating the ISG15 gene rescue is successful. As shown in Figure 5D, PRV-gE expression was much lower in ISG15^-/-^ cells transfected with ISG15AA-expressing plasmid than that of ISG15^-/-^ cells with empty vector. In line with the above results, ISG15 silencing impairs anti-PRV response of IFN*α* via a monomer form.

**FIG 5.**
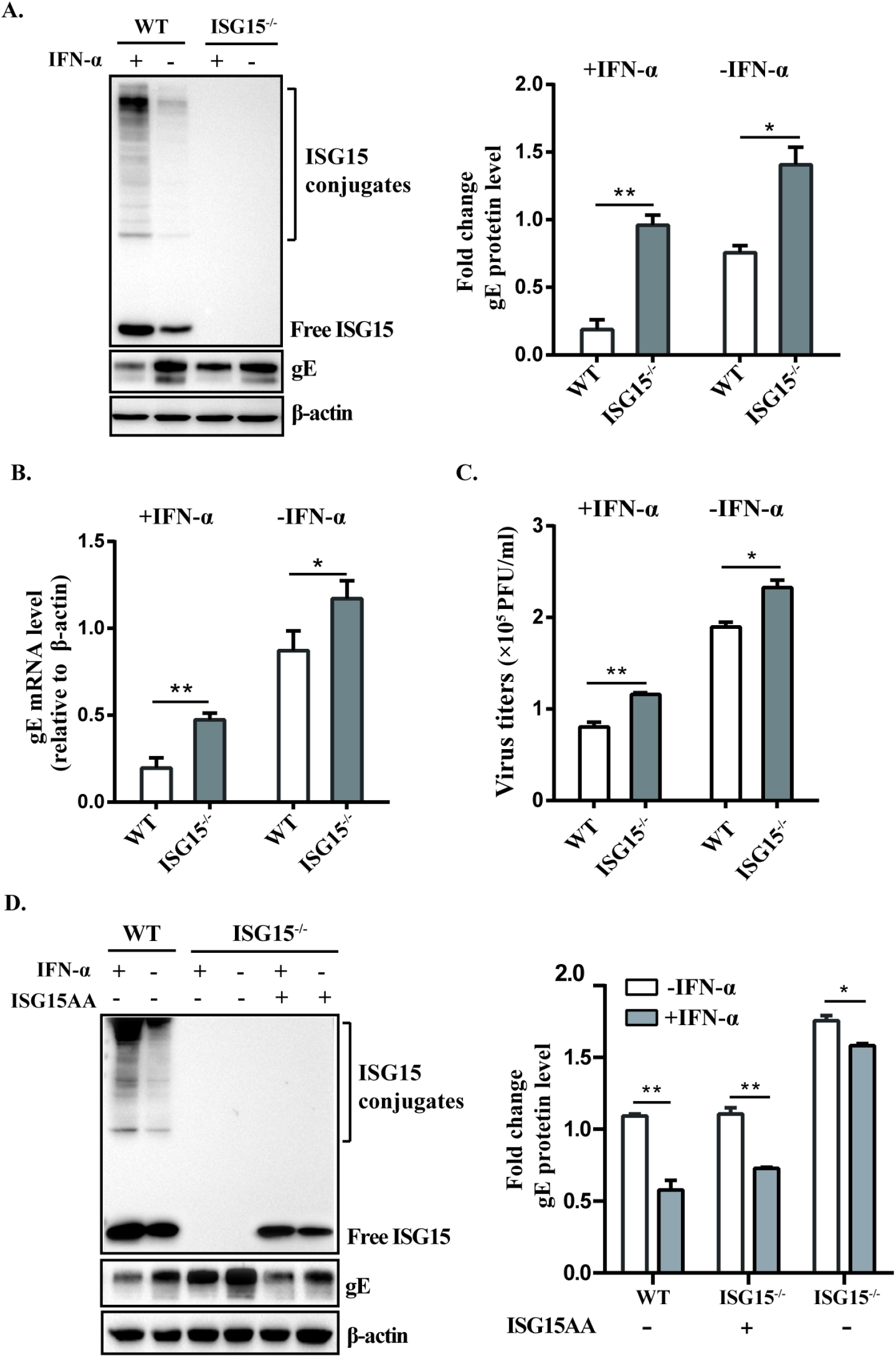
ISG15 promotes IFNα antiviral activity against PRV. (A to C) Prior to infection with PRV at an MOI of 1, WT and ISG15^-/-^ cells were either treated with IFNα (1000 U/ml) or left untreated. Protein extracts from the cells were harvested 24 hpi and analyzed by Western blot. Relative fold changes in PRV-gE expression levels are indicated between WT and ISG15^-/-^ cells in the absence and presence of IFNα (B to C). Total RNA and supernatants were harvested 24 hpi to determine the PRV-gE mRNA level and virus titers by RT-qPCR and plaque assay, respectively. (D) ISG15^-/-^ cells were transfected with either a plasmid expressing ISG15 mutant or an empty plasmid. After 12 h, WT and ISG15^-/-^ cells were treated with or without IFNα (1000 U/ml) before infecting with PRV. Western blot was used to analyze the protein expression of ISG15 and gE,The data are presented as means standard error of the mean ± (SEM) of at least three independent experiments. *, *p*≤0.05 (*t*-test).

Altogether, our data supports that free ISG15 promotes IFN*α*-mediated antiviral activity against PRV.

### ISG15 facilitates the phosphorylation of STAT1 and STAT2

To understand the mechanism by which ISG15 participates in IFN*α*-mediated antiviral activity, we monitored the phosphorylation levels of STAT1 and STAT2 in WT versus ISG15^-/-^ cells following PRV infection with or without IFN-*α* treatment. As illustrated in Figure 6A and B, IFN*α* stimulation accelerated pSTAT1 and pSTAT2, while ISG15 silencing significantly inhibited the expression of pSTAT1 (Fig. 6A) and pSTAT2 (Fig. 6B) regardless of the inclusion or exclusion of IFN*α* treatment. Subsequently, subcellular fractionation and Western blot analysis show that the IFN*α*- induced nuclear translocation of STAT1/STAT2 in the ISG15^-/-^ cells was reduced accordingly (Fig. 6C). This observation was consistent with the data from immunofluorescence images (Fig. 6D), indicating that ISG15 silencing blocks the nuclear translocation of the STAT1 and STAT2. This provides strong evidence that ISG15 silencing inhibits STAT1 and STAT2 nuclear accumulation by blocking nuclear import.

**FIG 6.**
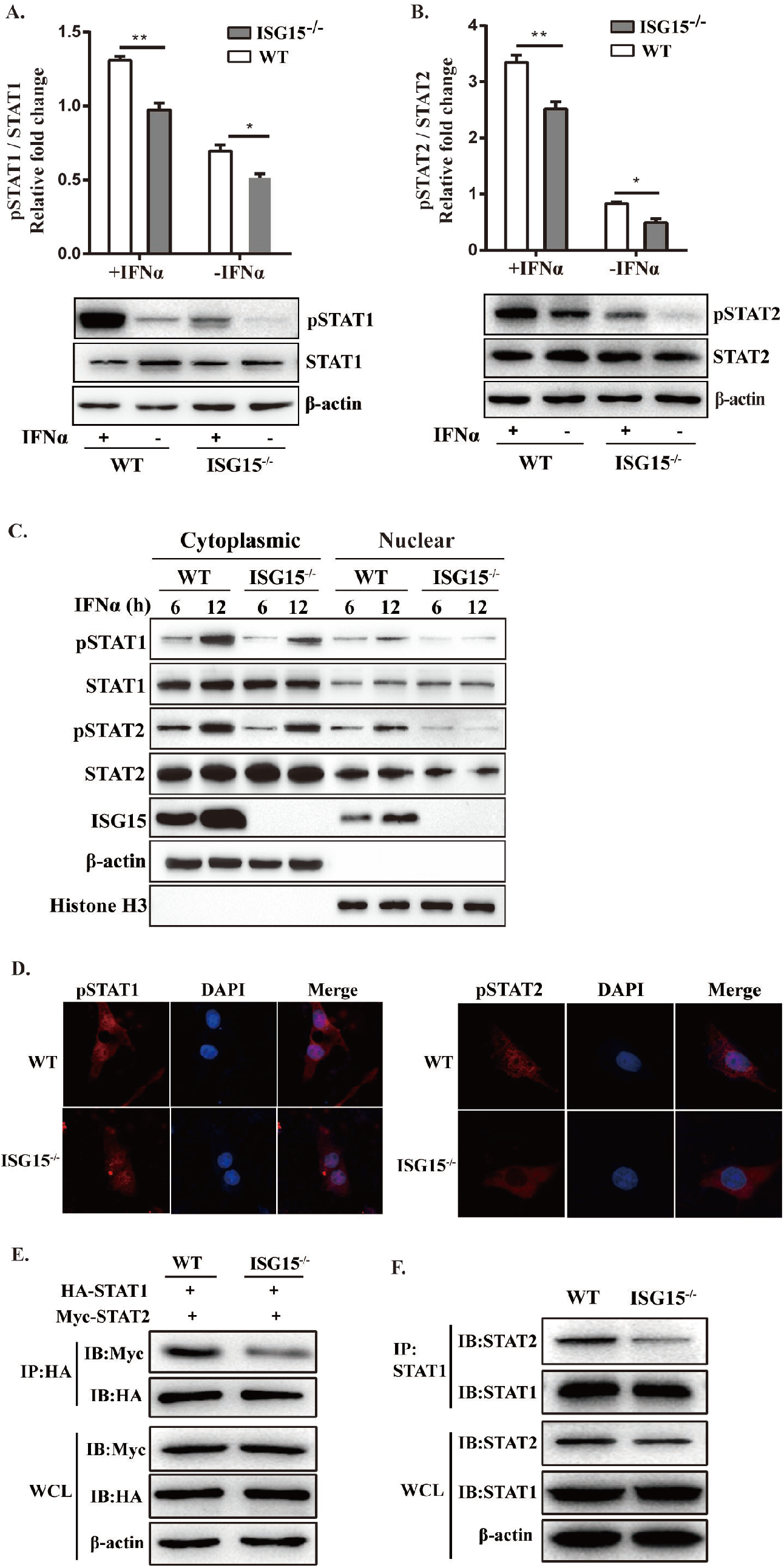
ISG15 facilitates the phosphorylation of STAT1/STAT2. (A to B) WT and ISG15^-/-^ PK15 cells were treated or un-treated with IFNα (1000 U /ml) and then infected 12 h later with PRV (MOI=1). At 24 hpi, the relative pSTAT1 and pSTAT2 levels were analyzed by Western blotting with the specified antibodies. The ratio of pSTAT1 (pSTAT2)and STAT1-tot (pSTAT2-tot) protein levels relative to control was calculated by ImageJ software. (C) Subcellular fractionation and Western blotting were used to identify phosphorylated STAT1 and STAT2 in nuclear and cytoplasmic fractions of WT and ISG15^-/-^ cells. The same blot was incubated with antibodies against *β*-actin and histone H3 as controls for loading and fractionation. (D) Immunofluorescence of WT and ISG15-/- cells transfected with HA-STAT1 or Myc-STAT2 and stimulated with IFNα. Nuclei (blue) and pSTAT1/pSTAT2 (red) were detected. (E) WT cells and ISG15^-/-^ PK15 cells were co-transfected with HA-STAT1 and Myc-STAT2. At 24 hpt, the cells were treated with IFNα and infected with PRV (MOI=1), and the STAT1 and STAT2 expression were analyzed by anti-HA and anti-Myc antibodies. (F) Endogenous STAT1 and STAT2 expression were detected from cells treated with IFNα and infected with PRV, with anti-STAT1 (Tyr 701) and anti-STAT2 (Tyr 690) specific antibodies. *, *p*≤0.05 (*t*-test).

ISG15 was found to be predominantly localized in the cytoplasm (Fig. 6C), the mechanism for ISG15 to influence this step is likely through altering the expression levels of the components essential for IFN signaling pathway. Since ISG15 silencing inhibited the phosphorylation of STAT1 and STAT2, we hypothesized that ISG15 deletion blocked the interaction of the STAT1 and STAT2. Thus, a co-immunoprecipitation assay (Co-IP) was performed to detect whether ISG15 impacted the interactions of STAT1 and STAT2. First, WT and ISG15^-/-^ cells were co-transfected with HA-STAT1 and Myc-STAT2. Twenty-four hours later, the cell lysates were immunoprecipitated with an anti-HA antibody and then immunoblotting with anti-Myc antibody, and the results showed the expression level of STAT1 in the presence of STAT2 in the samples from the two group cells (Fig.6E). In parallel, the presence of STAT1 in both samples was verified by IP with a Myc-tag antibody followed by blotting with an anti-HA-tag antibody. Meanwhile, the interaction between endogenous STAT1 and STAT2 was also analyzed by Co-IP assay, and similar results were observed (Fig. 6F). This indicated that ISG15 silencing attenuated the interaction between STAT1 and STAT2.

Collectively, these results suggest that ISG15 promotes the IFN*α*-induced phosphorylation of STAT1 and STAT2.

### ISG15 silencing inhibits ISGF3-induced ISRE reporter activity

The heterodimerization of phosphorylated STAT1 and STAT2 associates with IRF9, forming the ISGF3 complex that subsequently enters the nucleus to activate ISRE-dependent transcription [32]. We have found ISG15 silencing suppresses the phosphorylation of STAT1 and STAT2, so we speculate ISG15 may affect the formation of ISGF3 complex, thereby inhibiting the expression of ISGs. To dissect the idea that ISG15 is involved in the ISGF3 formation, WT or ISG15^-/-^ cells were treated with IFN*α* or left untreated following PRV infection. Subcellular fractionation and subsequent Western blot analysis showed that ISG15 silencing downregulated the phosphorylation level of STAT1 and STAT2 induced by IFN*α*, while had only slight impact on the expression of IRF9 (Fig.7A). This suggests a complete loss of ISG15 attenuates the form of ISGF3 complex, thereby inhibiting the ISGF3 translocation into the nucleus.

To study the effect of ISG15 on ISGF3-induced ISRE promoter activation, WT and ISG15^-/-^ cells were co-transfected with the plasmid-encoded STAT1, STAT2 and/or IRF9, together with ISRE-luciferase and Renilla luciferase reporters. Twelve hours later, the cells were infected with PRV and then collected for analysis of luciferase activity induced by ISRE. The result showed that ISRE promoter activity markedly diminished compared to the other control groups (p<0.05; Fig.7B), suggesting that ISG15 promoted to the ISRE promoter activation. Additionally, activated ISGF3 drives the transcription of ISGs, which are important for the control of viral infections [33]. We also detected the expression of IFIT1, 2’,5’-oligoadenylate synthetase 1 (OAS1) and myxovirus resistance protein A (MxA), the downstream transcription factors of IFN signaling. Results indicated that ISG15 deletion significantly down-regulated the transcription levels of IFIT1, MxA and OAS1 (Fig. 7C).

**FIG 7.**
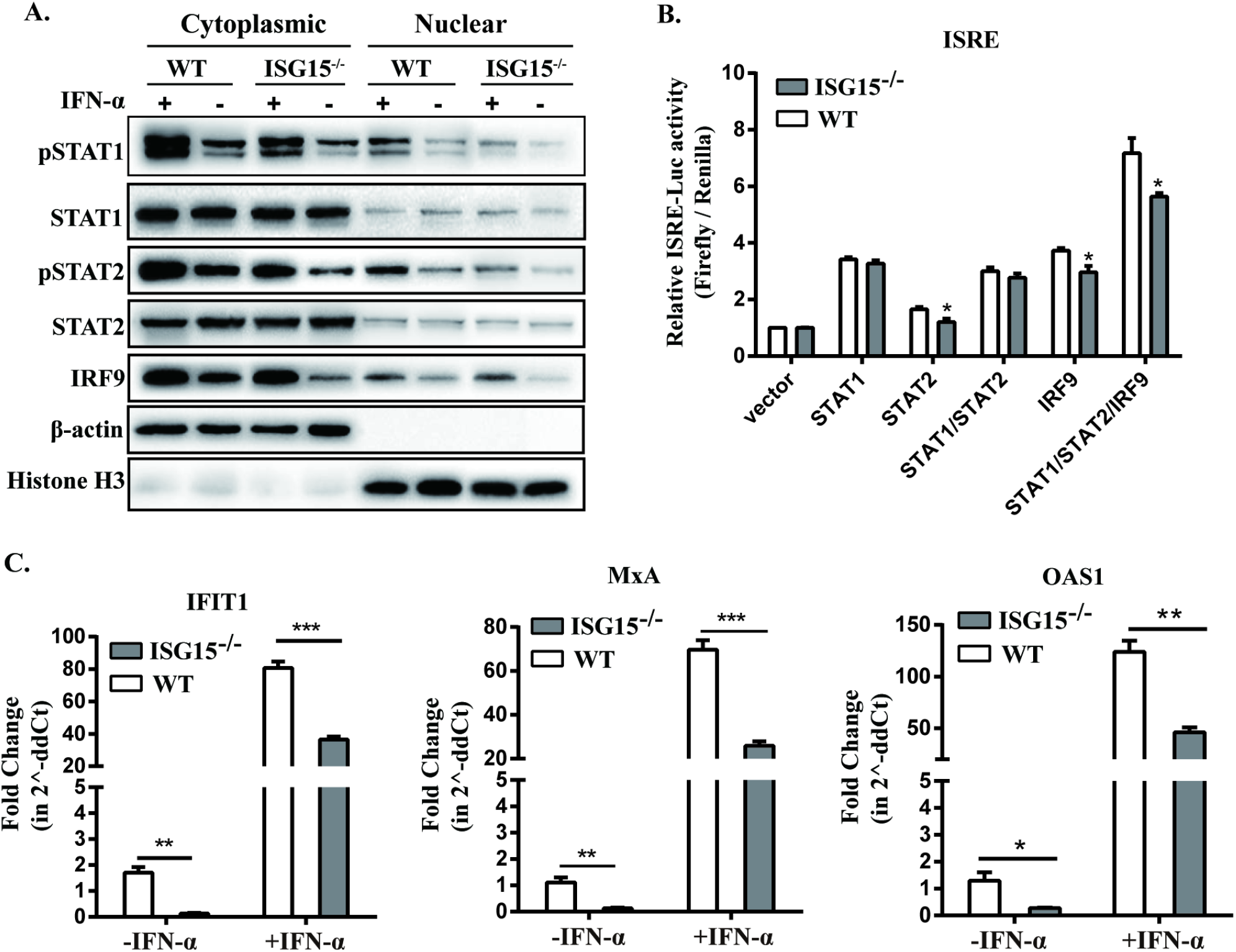
ISG15 silencing facilitates the formation of ISGF3. (A) WT and ISG15^-/-^ cells were treated or un-treated with IFN*α* and then infected with PRV. Subcellular fractionation and sequent Western blotting were used to detect p-STAT1, p-STAT2, total STAT1, total STAT2, IRF9 in the cytoplasmic and nuclear fractions of WT and ISG15^-/-^ cells. The same blot was incubated with antibodies against *β*-actin and histone H3 as controls for loading and fractionation. (B) WT and ISG15^-/-^ cells were co-transfected with STAT1, and/or STAT2, and/or IRF9, pGL4.17-ISRE-Luc (firefly luciferase) and pRL-TK (Renilla luciferase). ISRE promoter activity was measured at 24 hpi for Dual-luciferase reporter gene assay. (C) IFIT1, MxA and OAS1 mRNA levels were quantified the samples from WT or ISG15^-/-^ cells with or without IFNα treatment. Fold change in mRNA levels relative to the untreated group was calculated using the 2^△△ *CT*^ method, and the *β*-actin gene was used as the housekeeping gene. **, *p*≤0.001; *** *p*≤0.0001 (*t*-test).

Taken together, these data provided evidence that a complete loss of ISG15 blocked the ISGF3 formation, attenuated ISRE promoter activity and downregulated the transcription levels of ISGs.

### ISG15^-/-^ mice are more sensitive to PRV infection

Further, we examined the role of ISG15 in host defense to control PRV infection using genetically knockout ISG15 (ISG15^-/-^) mice. A survival analysis was first performed to confirm the role of ISG15 in host survival during PRV infection. C57BL/6N (B6, WT) and ISG15^-/-^ mice were infected with PRV-QXX via subcutaneous injection and monitored for 6 days post infection (dpi). The infected ISG15^-/-^ mice began to die at 3 dpi with severe itchiness symptoms such as. The survival rate of ISG15^-/-^ mice reached 100% until 6 dpi, while the survival rate of WT mice was only 46.7% (Fig. 8A), suggesting that the occurrence of increased susceptibility to PRV infection in the complete loss of ISG15. We also observed a marked increase in ISG15 protein in brains of infected WT mice (Fig. 8B), which is consistent with the results obtained in the cell model. PRV-gE gene copies and viral titers of brains were examined by RT-qPCR and plaque assay individually for each mouse. As expected, PRV-gE gene copies and viral titers in ISG15^-/-^ mice were significantly higher than those in WT mice (Fig. 8C to D).

**FIG 8.**
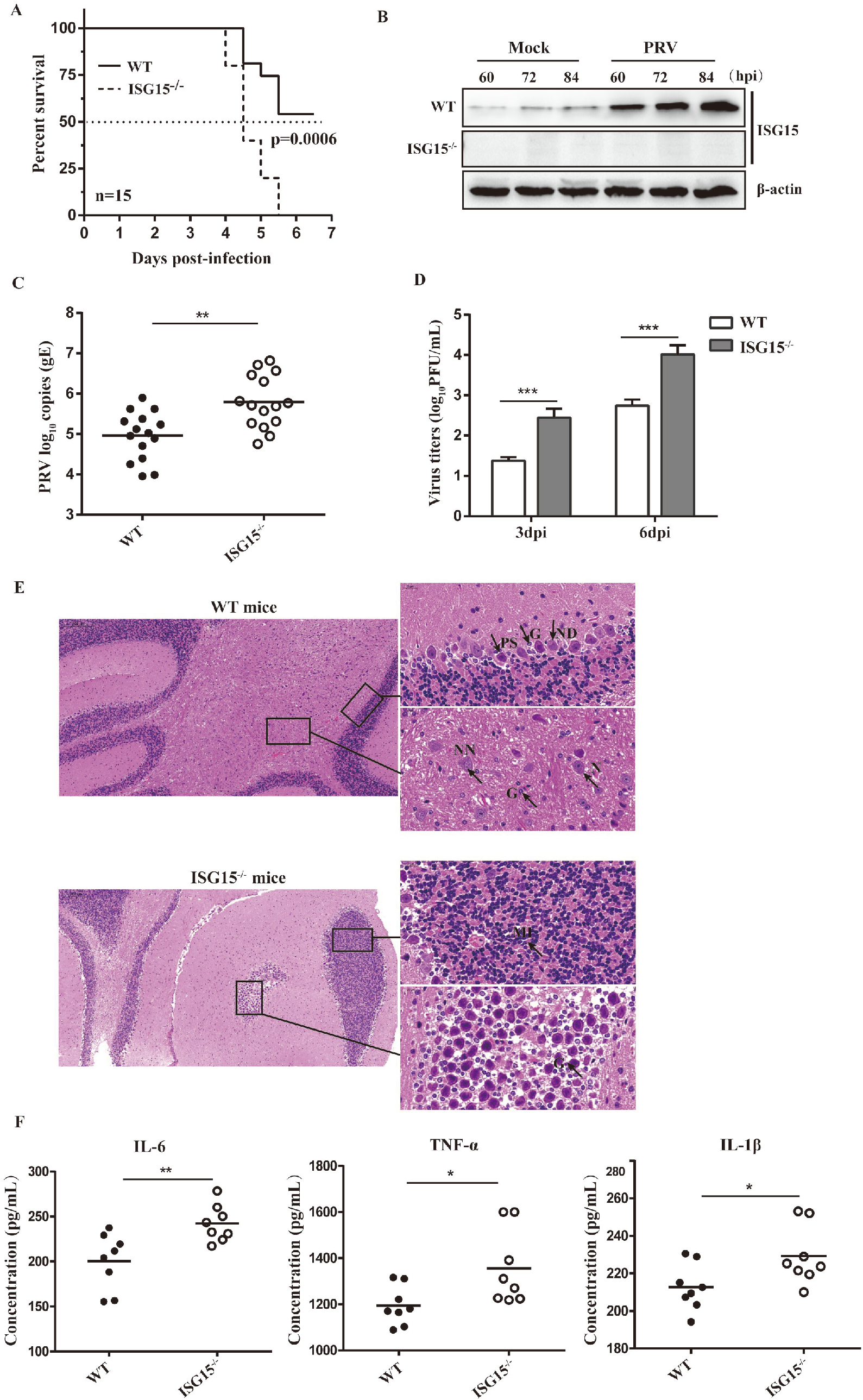
PRV challenge assay in vivo. Seven-week-old male ISG15^-/-^ mice (n=15) and WT mice (n=15) were inoculated with 5×10^3^ TCID_50_ of PRV-QXX subcutaneous. (A) Survival of the infected WT and ISG15^-/-^ mice was monitored until day 6 after infection. Statistical significance was determined by the log-rank test. (B to D) Brain tissues of the infected mice were collected at different days post-infection. The ISG15 expression, PRV-gE copies and viral titers were detected by Western blotting, RT-qPCR, and plaque assay respectively. (E) The histopathological features of brains of the infected WT and ISG15^-/-^ mice. The brain tissues were sectioned and stained with hematoxylin-eosin. Magnified images of the regions with black rectangles in infected WT and ISG15^-/-^ mice, respectively. Representative images are shown: ND: nuclear disintegration of Purkinje cell; N: neurons; NN: necrotic neurons; G: glial cells; PS: shrinkage of Purkinje cells; P: Purkinje cells; PN: necrotic Purkinje cells; MI: mononuclear cellular infiltration. (F) The concentrations of IL-6, TNF-α and IL-1β in serum of infected WT and ISG15^-/-^ mice were determined by ELISA.

**FIG 9.**
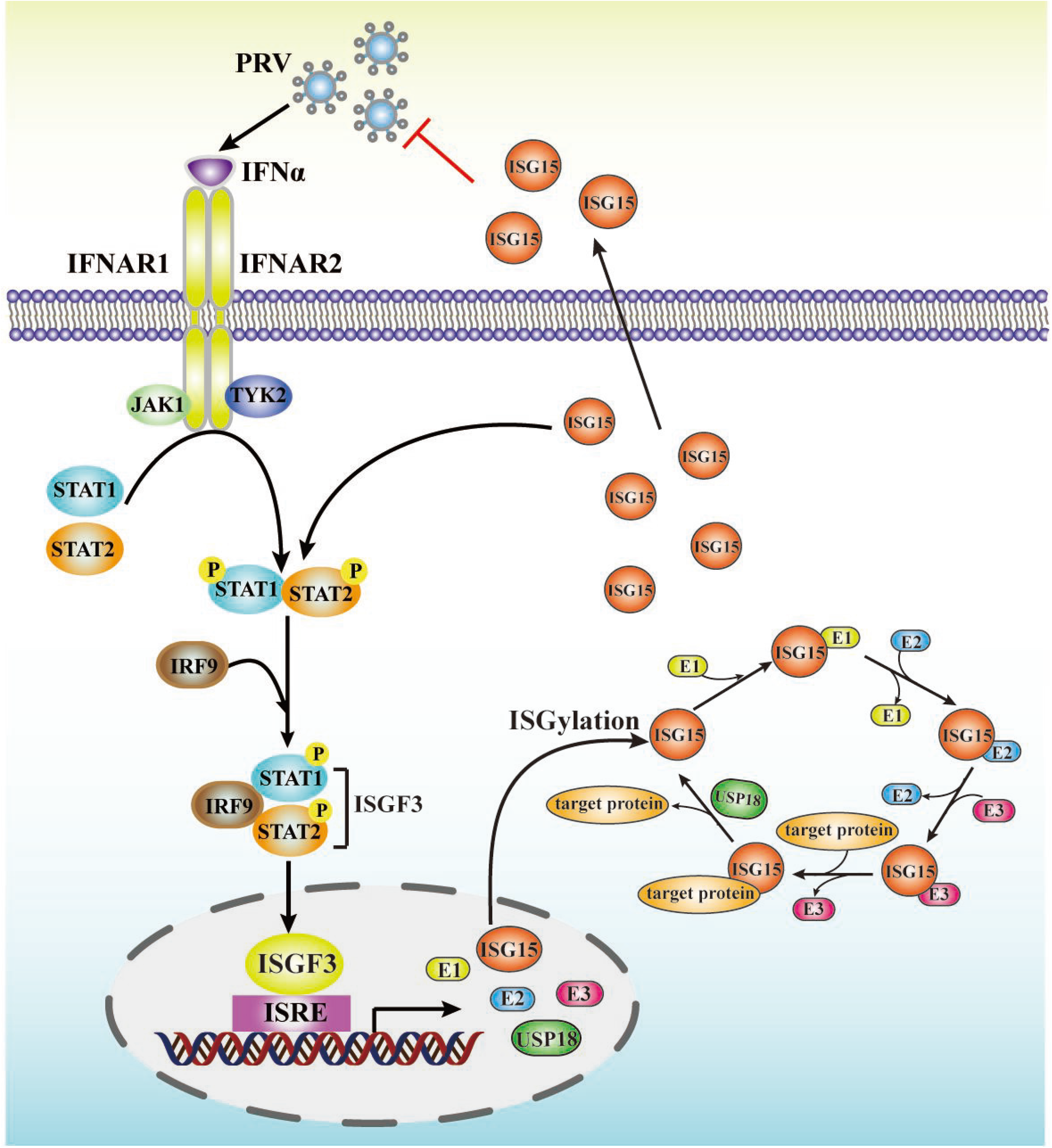
Mechanism that has been provided. Type I IFN induced by PRV infection mediates antiviral response by upregulating ISG15. Our work demonstrates that increased ISG15 positively regulates IFN*α*-induced antiviral activity against PRV, by facilitating phosphorylation of STAT1 and STAT2. This regulation results in increased ISG15 inhibiting PRV replication (red line).

Encephalitis caused by PRV infection in the central neuron system is a pivotal factor leading to animal death [34]. Therefore, we detect the degree of encephalitis in the brain tissues of infected WT and ISG15^-/-^ mice by histopathological observation. Hematoxylin-and-eosin staining showed greater inflammatory damage, necrotic neurons, and more glial cells in ISG15^-/-^ mice than those in WT mice (Fig. 8E). We further determined IL-6, TNF-*α* and IL-1β protein levels in the brains of the WT and ISG15^-/-^ mice by ELISA. Increased protein levels of IL-6, TNF-*α* and IL-1β were observed in ISG15^-/-^ mice compared to WT mice (Fig. 8F). This suggests that ISG15 may have a direct or indirect role in encephalitis.

Taken together, our observations imply that ISG15 plays a critical role in host anti-PRV response.

## Discussion

To date, the PRV-host interactions that induces ISG15 expression and the impact of the ISG15 monomer and conjugates on PRV replication remained unexplored. Here, we establish that although the ISG15 abundance are triggered by PRV infection, they are subsequently tempered, but not completely abrogated by viral gene expression. Preventing PRV gene expression and viral growth greatly reduced ISG15 accumulation. Therefore, both free and conjugated ISG15 accumulation in response to PRV infection was dependent on viral gene expression and viral growth. Moreover, deletion of ISG15 remarkably promoted PRV replication, indicating that ISG15 has a significant anti-PRV function. Additionally, we noticed that ISG15 had to accumulate in large amounts before virus infection to carry out its anti-PRV role. ISG15 seems to affect a stage in the PRV cycle before virus release, since PRV titers increased in both cell-associated and released virus following ISG15 silencing (Fig. 3). In contrast, it has been recently reported that ISG15 blocks the entry and/or uncoating phase of the murine norovirus life cycle [24]. Furthermore, ISG15-deficient mice display more sensitive to PRV infection, indicating ISG15 exerts a critical role for restricting PRV infection against PRV *in vitro* and *in vivo*.

We found that ISG15 exerts its anti-PRV effect relying on ISG15 monomer, as demonstrated by the effect of an unconjugated form of ISG15, the inhibition of ISGylaiton by UbE1L silencing, or the enhancement of ISGylation by USP18 silencing. It is possible that extracellular ISG15 monomer directly interferes with PRV replication to prevent viral infection, as evidenced by the high levels of ISG15 expression before PRV-gE expression (Fig.1). In other words, when ISG15 is expressed at high levels before virus infection, as in cells stimulated by IFN*α*, no viral proteins are present to counteract the ISG15 antiviral activity. Another possibility is that ISG15 accumulation may promote type I IFN signaling and/or the expressions of ISGs induced by viral infection. The positive correlation between PRV infection and ISG15 expression was also observed from the infected mice (Fig.8B and 8D), pointing to the antiviral role of ISG15 in PRV infection *in vivo*. Our result contrast with the previous studies that ISG15-deficient patients who display no enhanced susceptibility to viruses *in vivo*. This reflects the ISG15 function may vary depending on the virus and host species.

Type I IFN is critical for controlling PRV infection *in vitro* and *in vivo*. Recent investigations demonstrated that ISG15 acted as a negative regulator of type I IFN signaling exerted antiviral response during viral infection [31, 33]. However, our results provided some evidence supporting ISG15 as a positive regulator of IFN*α*- mediated antiviral response against PRV as following: 1) ISG15 silencing impairs the antiviral activity of IFN*α* against PRV; 2) ISG15 deletion blocks STAT1 and STAT2 phosphorylation through inhibition of the interaction between STAT1 and STAT2; 3) ISG15 facilitates the ISGF3 complex formation and ISRE promoter activity; and 4) The transcription level of ISGs genes induced by IFN*α* greatly reduced in ISG15 knockout cells. These data demonstrate ISG15 as a key positive regulator in IFN signaling and confirm its importance in host defense response against PRV infection, which may be more broadly against other viruses as well.

We found that ISG15 was involved in two crucial steps in IFN signaling, including the active pSTAT1 and pSTAT2 and the ISGF3 formation (Fig. 7 to 8). This may be partly because ISG15 knockout reduced interactions of STAT1 and STAT2 (Fig. 6E to F). Since ISG15 mainly localizes in the cytoplasm, we suppose the mechanism for ISG15 to impact this step is likely direct. The result that the lack of ISG15 decreases STAT1 and STAT2 phosphorylation with hindering interactions between STAT1 and STAT2 suggests that ISG15 may be involved in the formation of the STAT1-STAT2 heterodimer. Although the regulation of the ISGF3-mediated transcription of ISGs in the nucleus in not well understood, we demonstrate that ISG15 carries out a critical positive regulator in this process. ISG15 seems to function by enhancing the ISGF3 recruitment to the promoter of ISGs and promoting the transcription of ISGs, this is complex. ISGF3 complex is essential for ISRE activation. Thus, we speculate that ISG15 may act as a regulator promoting ISGF3 to its ISGs promoters for efficient gene transcription. A similar mode of action is also observed in Bclaf1 that regulated the type I interferon responses and was degraded by alphaherpesvirus US3 [35].

We further studied the effects of ISG15 on PRV infection *in vivo* by using ISG15^−/−^ mice model. The results identified that ISG15^−/−^ mice displayed increased morbidity and mortality rates, viral replication, as well as promotes development of viral encephalitis in the brains of mice. This finding confirms that ISG15 positively regulates host anti-PRV effect *in vivo*, which may be broadly for other viruses as well. Others and our studies have highlighted a critical role of ISG15 during viral infections, and it could be a viable option for developing therapeutic target for controlling PRV.

## MATERIALS AND METHODS

### Cell culture and virus

Porcine kidney epithelial cells (PK15) were cultured at 37°C in 5% CO_2_ in Dulbecco’s modified Eagle medium (DMEM; Gibco, Grand Island, NY, USA) supplemented with 10% fetal bovine serum (FBS; Gibco) and 1% penicillin-streptomycin (DingGuo, Beijing, China). The PRV-QXX virus was preserved in our laboratory. For experiments, PRV was amplified in PK15 cells, and virus titers were determined using a plaque assay, as previously described [21]. PRV infections was performed at an MOI of 1 PFU/cell.

### Chemicals and chemical treatments

IFNα (Pbl, NJ, USA) was dissolved in 0.1% BSA and used at a final concentration of 1000 U/mL. The viral DNA polymerase inhibitor phosphonoacetic acid (PAA) was dissolved in deionized water and utilized at a concentration of 300 *μ*g/mL (GlpBio, Montclair, CA, USA). The chemicals was added to the cultures at the indicated concentrations.

### Quantitative RT-PCR and Western blots

According to the protocol of the manufacturer, RNA was extracted from cells using the TRIzol reagent (Takara, Shiga, Japan) and reverse transcribed using the PrimeScript™ RT reagent Kit (Takara). Quantitative RT-PCR was used to determine gene expression using the SYBR Green Realtime Master Mix (Takara, DaLian, China). Table 1 contains a list of all primers used in this study. All values were normalized to the level of *β*-actin mRNA, and relative expression was calculated using the comparative cycle threshold (2^-ΔΔCT^) method.

The cells were harvested and washed twice with PBS before being lysed with RIPA. After 15 minutes of centrifugation at 13,000 rpm, the supernatant fraction was collected. The BCA Protein Assay Kit was used to assess the protein concentration in supernatants (Beyotime Biotechnology, Shanghai, China). Equivalent quantities of each protein sample were electrophoresed on SDS-PAGE gels and transferred to PVDF membranes (Pall Corporation, Ann Arbor, MI, USA). The primary antibodies directed against the following proteins were: anti-ISG15 (1:3000 dilution; Abcam); anti-*β*-actin, anti-HA, anti-myc (1:3000 dilution; Proteintech, Wuhan, China); anti-PRV-glycoprotein E (gE); anti-phospho-Tyr701 STAT1; anti-STAT1 (1:3000 dilution; Cell Signaling); anti-phospho-Tyr690 STAT2; anti-STAT2 (1:3000 dilution; Cell Signaling); anti-IRF9 (1:3000 dilution; Cell Signaling); anti-USP18 (1:3000 dilution; Cusabio Wuhan, China). Secondary antibodies conjugated with horseradish peroxidase against rabbit or mouse (1:5000 dilution; Santa Cruz) were used. The ECL Western blotting Analysis System was used to reveal protein bands (Millipore, United States). Densitometry was performed with ImageJ software and standardized against *β*-actin.

### Immunofluorescence assay

PK15 cells were plated into a confocal dish and transfected with HA-STAT1 or myc-STAT2 plasmid. 4% paraformaldehyde was used to fix the monolayer cells, and 0.5% Triton X-100 were used to permeabilized at 4°C with (Solarbio Life Science, Beijing, China). Following a wash with PBS, cells were permeabilized in blocking solution (5% bovine serum albumin in PBS) for 1 h. Fixed cells were treated with a primary antibody specific for PRV-gE followed by an Alexa Fluor 488-conjugated secondary antibody against mouse (Proteintech, Wuhan, China). 4’, 6-diamidino-2-phenylindole (DAPI) was used to stain the cell nuclei (Solarbio). Fluorescence pictures were acquired by confocal laser scanning microscopy (Olympus, Tokyo, Japan).

### ISG15 mutant plasmid

The nonconjugative ISG15 plasmid pCAGGS-ISG15AA was constructed from pCAGGS-ISG15 using the site-directed mutagenesis kit (Beyotime), with the following primer pair: forward, 5’- TATA TGAATCTGCGCCTGCGGGCGGCCGGGACAGGG-3’, and reverse, 5’- CCCTGTCCCGGCCGCCCGCAGGCGCAGATTCATATA-3’.

### siRNA silencing

Twenty-four hours prior to transfection, PK15 cells were plated in 24-well plates. The cells were transfected with control small interfering RNAs (siRNAs), or specific siRNAs against UbE1L or USP18 using with 1 μL Lipofectamine RNAiMAx reagent (Invitrogen) per well. At 24 hpt, the cells were infected with PRV (MOI=1). At four hpt, the culture media was changed with fresh medium containing 1000 U/mL IFNα, which was maintained throughout the infection duration. At 24 hpi, supernatants were collected for the viral titration, and cells were extracted for Western blotting and RT-qPCR analysis. The siRNAs sequences employed in this study were as follows: UbE1L no. 1: GCACUUCCCACCUGAUAAA; UbE1L no. 2: CAGCC UCACUCUUCAUGAU; USP18 no. 1: GUCUCCAGAAGUACAAUAUTT; USP18 no. 2: CCAGUGUACUUAUGGAAAU; NC: UUCUCCGAACGUGUCACGU.

### ISRE-luciferase reporter assay

Co-transfection of PK15 and ISG15^-/-^-PK15 cells with the identified plasmid and the ISRE-Luc reporter plasmid (100 ng) plus the internal control pRL-TK reporter plasmid was performed (5 ng). Cells were treated with IFNα (1000 U/mL) for 12 h and were harvested to conduct dual-luciferase reporter assay (Promega, Madison, WI). Firefly luciferase activity values were normalized to Renilla luciferase activity, and the relative fold changes in IFN-treated samples compared to IFN-untreated control were calculated.

### Co-immunoprecipitation (Co-IP) assays

PK15 cells were co-transfected with HA-STAT1 and myc-STAT2 plasmids and then lysed with ice-cold lysis buffer (25 mM Tris-HCl, pH 7.4, 1% NP-40, 150 mM NaCl, 1 mM EDTA) supplemented with protease inhibitors (Sigma). Cell lysate was cleared by centrifugation at 14,000×g for 5 min at 4°C. Primary antibodies against HA, STAT1 or STAT2 (dilution 1:1000; Proteintech) were added to the supernatants. After three washes with TBS, SDS-PAGE sample buffer was added, and proteins were separated by SDS-PAGE and immunoblotted to determine STAT1 and STAT2 interaction.

### PRV challenge assay *in vivo*

The ISG15 knockout mice were generated from the Cyagen Biosciences (Cyagen, China). The seven-week-old male ISG15^-/-^ mice and WT mice were randomly divided into two groups consisting of 15 mice each, respectively. The mice had free access to pelleted food and water during the experimental period. Each mouse was challenged by the subcutaneous infection with 50μl of DMEM containing 5×10^3^ TCID_50_ of PRV-QXX. All the mice were monitored daily, and the mortality was recorded from 1 to 6 days post-infection. The brain samples were excised to detect the viral titer and the gE gene copies by plaque and RT-qPCR, respectively. Blood serum were also collected and kept at 4 °C to detect inflammatory factor through specific antibodies by enzyme-linked immunosorbent assay (ELISA). In parallel, the brain tissues were fixed in neutral-buffered formalin for histological analysis. All the animal experiments used in this study were approved by the Animal Ethics Committee of Henan Agricultural University.

### Statistical analysis

GraphPad Prism 8 software was used to conduct statistical comparisons. The difference between groups was determined using Student’s *t*-tests, and *P* values less than 0.05 were considered statistically significant (p < 0.05). The standard errors of the mean (SEM) of at least three independent experiments are shown for each data.

## ACKNOWLEDGMENTS

This study was financially supported by grants from the National Natural Science Foundation of China (31902268 and 31772781).

